# MINTyper: An outbreak-detection method for accurate and rapid SNP typing of clonal clusters with noisy long reads

**DOI:** 10.1101/2020.05.28.121251

**Authors:** Malte B. Hallgren, Søren Overballe-Petersen, Ole Lund, Henrik Hasman, Philip T. L. C. Clausen

**Affiliations:** National Food Institute, Technical University of Denmark, Kemitorvet 204, 2800 Kgs. Lyngby, Denmark; Department of Bacteria, Parasites and Fungi, Statens Serum Institut, Copenhagen, Denmark

## Abstract

For detection of clonal outbreaks in clinical settings, we present a complete pipeline that generates a SNP-distance matrix from a set of sequencing reads. Importantly, the program is able to handle a separate mix of both short reads from the Illumina sequencing platforms and long reads from Oxford Nanopore Technologies’ (ONT) platforms as input.

MINTyper performs automated reference identification, alignment, alignment trimming, optional methylation masking and pairwise distance calculations. With this approach, we could rapidly and accurately cluster a set of DNA sequenced isolates, with a known epidemiological relationship to confirm the clustering. Functions were built to allow for both high-accuracy methylation-aware base-called MinION reads (hac_m Q10) and fast generated lower-quality reads (fast Q8) to be used, also in combination with Illumina data. With fast Q8 reads a higher number of base pairs were excluded from the calculated distance matrix, compared to the high-accuracy methylation-aware Q10 base-calling of ONT data. Nonetheless, when using different qualities of ONT data with corresponding input parameters, the clustering of isolates were nearly identical.

## INTRODUCTION

Until the 21st century the field of microbial diagnostics was dominated by non-computational methods. These methods ranged from cultivation and microscopic visualization to a wide variety of laboratory-based assay-technologies. Shared shortcomings of these methods were long diagnostic times and/or relatively low precision. Often the identity of a pathogenic isolate could not be determined with greater accuracy than the sample’s species or genus, and it could take several days, if not weeks, to perform additional tests (1). The introduction of DNA sequencing in the late 1970’s by Sanger, and the subsequent improvements of the concept in the form of 2nd and 3rd generation sequencing, has allowed for better and faster analysis of microbes at a phylogenetic level (2). Illumina 2nd generation sequencing technology have dominated the market for the previous 10 years, as it allows for precise and cost-effective sequencing, when a large pool of samples are sequenced together using multiplexing (3).

Due to the requirement of sample multiplexing in order to make Illumina platforms cost-effective in a clinical setting, researchers are looking for more agile sequencing alternatives with a shorter turnaround time. The ONT MinION platform offers great potential due to the low cost of the machine, low average sequencing price per run and short turnaround time, thus allowing for much smaller pools of samples to be sequenced.

One of the most significant factors that currently is preventing 3rd-generation long-read sequencing-platforms from replacing the short-read sequencers is the increased error rate of long-read sequencing-technologies (4). For some research purposes an increased error rate can be overcome, but especially when working with genetics and phylogeny, where the difference between defining if an isolate is part of an outbreak or not may be down to a few single nucleotide polymorphisms (SNPs), error-prone sequences can distort the analysis.

When screening for clonal bacterial outbreaks a widely used method is to perform a SNP-typing analysis (5). Assuming that the SNPs are the result of random point-mutations, i.e. not a result of recombination, SNP distances between isolates can be used as measurements of relatedness.

Here we present for the first time an automated SNP-typing method (MINTyper), that based on long-read sequencing can infer the same clonal clusters as methods based on short-read sequencing. The method was validated on a test set containing 12 *Escherichia coli* isolates separated into two subgroups: 1) Outbreak isolates of ST410 type (n=6) and 2) non-Outbreak isolates of ST410 type (n=6) based on their epidemiological relationship (patients travel and hospitalization data). Furthermore, MINTyper has been designed to handle a mix of short-read and long-read sequencing samples. This enables comparison of historical data from older sequencing platforms with new data from long read sequencing platforms.

## MATERIALS AND METHODS

### Complete pipeline

MINTyper is a complete pipeline to identify and cluster clonal bacterial outbreak strains, including automatic identification of a bacterial reference sequence. The pipeline is composed of five main steps: Identification of reference sequence (if one is not provided by the user), reference guided multiple alignment, trimming of alignments, SNP calling and finally clustering the sequences using Neighbor-Joining or infer phylogenetic relationships using maximum likelihood. The pipeline is freely available as open-source at: https://github.com/MBHallgren/MINTyper and as web-service at: https://cge.cbs.dtu.dk/services/MINTyper.

### Data

The data used to test the performance of MINTyper originates from 12 *E. coli* isolates that has been sequenced by Statens Serum Institut (SSI) in Denmark. The 12 isolates had previously been studied using Illumina sequencing to perform Multi-Locus Sequence Typing (MLST) and core-genome MLST (cgMLST) analysis (6). It was found that all of the isolates were of the sequence type 410 (ST410) (6). Six isolates (Ec01-Ec06) were all of the same bacterial clone (cgMLST type CT587) from patients who had been hospitalized in Denmark concurrently at the same ward and thus were part of the same outbreak. The remaining six isolates (Ec07-Ec12) were acquired from patients, who had never been hospitalized with any of the 11 other patients in this study. Rather, these six patients had previously been hospitalized outside of Denmark and after returning to Denmark for further treatment had been found to be colonized or infected with isolates belonging to six non-related cgMLST types. This knowledge of the phylogenetic relationship of the isolates will be used to benchmark the quality of the final distance matrix calculated by MINTyper.

Each bacterial isolate was sequenced using both Illumina’s MiSeq sequencer and Oxford Nanopore’s MinION MK1B sequencer. For Illumina sequencing the DNA was extracted using the QIAGEN DNeasy Blood & Tissue kit and the library was prepared with the Nextera XT kit. Afterwards, sequencing adaptors were removed and reads end-trimmed to quality Q ≥ 20 using Trimmomatic v0.36 (7). For ONT sequencing, the DNA was extracted with the Beckman Coulter GenFind v2 kit with a DynaMag-2 magnet. The library was prepared with the 1D ligation barcoding protocol followed by sequencing with a R9.4.1 flow cell. Base-calling, demultiplexing and conversion to fastq format from the raw fast5 reads were done using Albacore v2.3.4. Sequencing adapters were removed with Porechop v0.2.3 (8). Finally, quality filtering to Q ≥ 8 were done using NanoFilt (9). For the high-accuracy Q10 ONT reads the base-calling was performed with Guppy 3.6.0 with high-accuracy methylation-aware configuration.

### Identification of reference sequence

The Center for Genomic Epidemiology provides a variety of sequence databases that are pre-indexed for use with KMA alignment and are automatically updated weekly by pulling changes from NCBI (10). From these, a database containing a total of 20,377 complete bacterial chromosomal reference sequences, excluding plasmid sequences, was downloaded from: http://www.cbs.dtu.dk/public/CGE/databases/KmerFinder/ version/20201028/bacteria.tar.gz. This database was searched using KMA v1.3.8 (11) with the “-Sparse” option, which identifies the reference in the database with the largest *k*-mer overlap to a set of query sequences.

### Reference-guided multiple alignment

Reads were aligned to the reference using KMA v1.3.8 (11), with the preset option “-mint2” for the Illumina sequences and “-mint3” for the ONT sequences. In addition to aligning the reads, KMA produces a consensus sequence for each sample, where single positions are signified with upper-case bases at significant positions and lower-case bases at positions that did not fulfill the SNP-calling criteria. These criteria are for the “-mint2” preset option: reads are mapped unambiguously and positions have a depth of at least 10, at least 90% support and are significantly over represented using a McNemar test with an *α* of 0.05. Whereas for the “-mint3” preset option these criteria are: reads are mapped unambiguously and positions have a depth of at least 10, at least 70% support and are significantly over represented using a McNemar test with an *α* of 0.05.

### Trimming of alignments

Trimming multiple alignments have proven useful when determining the phylogeny between closely related sequences. Highly divergent areas are removed in order to select the conserved blocks determining the clonal relationship between the isolates (12)(13). This ensures a phylogeny based on vertical evolution rather than horizontal evolution, where entire blocks of sequences are inherited in a single evolutionary event. In addition, this eliminates genomic regions with low sequencing quality. For outbreak investigation it is important to ensure a high specificity over sensitivity in order to distinguish whether a strain is part of an outbreak or not. Therefore core-genome SNPs are identified after trimming of alignments (14)(15). CCPhylo v0.2.2 was developed to trim the alignments across all isolates to only include positions that were significant according to the SNP calling criteria. Additionally, SNPs located within a proximity of 10 bases of each other were trimmed away, to reduce the effect of hyper-variable regions leading to sub-optimal alignments and remove regions that likely originated from horizontal evolution (12) (13) (16) (17). Additionally, DCM-methylation motifs (CCWGG) were trimmed away from the fast Q8 ONT data, which was base-called with Albacore, as these motifs constitute as much as 95% of discrepancies between Illumina and ONT data according to David R. Grieg et al. 2019 (18).

### Clustering and phylogenetic analysis

CCPhylo v0.2.2 was used to identify SNP differences between the isolates, and perform a subsequent hierarchical clustering using the Neighbor-Joining algorithm (19). As an alternative to Neighbor-Joining, maximum likelihood trees were generated using IQtree v2.0.3 (20) with parameters: “– seqtype DNA-seed 256”, and FasTree v2.1.11 (compiled with double precision) (21) with the parameters: “-gtr -nt”.

### Performance evaluation and comparison

MINTyper was evaluated on the data set containing 12 *E. coli*, where Illumina and ONT sequencined data were treated as separate samples to test the cross-platform performance of MINTyper. MINTyper was compared to MASH v2.2 (22) and CSIPhylogeny v1.4 (17) to evaluate the performance of MINTyper compared to other methods. The CPU-time and peak memory were measured for MASH, CSIPhylogeny and the individual parts of the MINTyper pipeline using GNU time v1.7.

## RESULTS AND DISCUSSION

### Automated reference detection

The best matching reference for the dataset of 12 ST410 *E. coli* was automatically identified by MINTyper as “Escherichia coli strain AMA1167 chromosome, complete genome”, both when searching the Illumina and ONT data as a combined dataset and individually. This result was anticipated, as this reference sequence is the published complete genome from the same danish outbreak that six of the input samples belong to (23).

### Trimming of alignments

The resulting output from running the MINTyper pipeline with the 12 ST410 *E. coli* isolates were a distance matrix and a Newick file for each of the three runs: One using fast Q8 ONT data with no trimming, one using fast Q8 ONT data with a minimum proximity of 10 bases between SNPs and DCM masking, and one using hac_m Q10 ONT data along with a minimum proximity of 10 between SNPs and no DCM masking). The ΔSNP between the Illumina reads and the ONT MinION reads can be seen in Table 1.

**Table 1.**
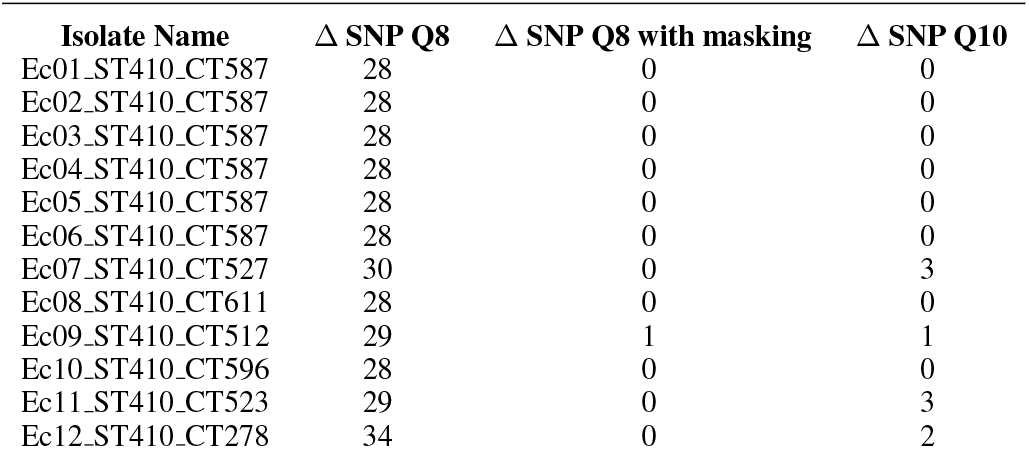
Overview of the number of SNPs differences between the consensus sequences generated by sequencing the same isolate on an Illumina platform and ONT MinION platform without trimming alignments and DCM methylation masking on fast Q8 data, alignment trimming and DCM methylation masking on fast Q8 data and alignment trimming but not DCM methylation masking on hac_m Q10 data. Alignment trimming were performed with a minimum distance of 10 between accepted SNPs.

The results of Table 1 shows that 11 out of 12 isolates had completely identical consensus sequences after DCM masking and alignment trimming was used with fast base-called Q8 data. Where Ec09 ST410 CT527 had a single SNP discrepancy.

As expected, the hac_m Q10 data no longer had most of the SNPs caused by the DCM sites. However, Ec07 ST410 CT512, Ec09 ST410 CT527, Ec11 ST410 CT527 and Ec12 ST410 CT278 had 1-3 SNP discrepancies each.

Distance matrices after alignment trimming with the proximity: 0, 10, 20, 50 and 100 between called SNPs have been included in S1 for the combination of Illumina with ONT fast Q8 or ONT hac_m Q10 respectively.

### Clustering and phylogenetic analysis

The three multiple alignments corresponding to the parameter setting described in table 1 were clustered using Neighbor-Joining, and were visualized using FigTree v1.4.4. Without alignment trimming it is clear that the systematic errors from the different technologies is stronger than the true relationships between the samples (See Figure 1). Trimming the alignments of Illumina and fast Q8 ONT data, with a minimum distance between accepted SNPs of 10, and masking out DCM methylation-sites revealed the true clustering (see Figure 2). Each isolate, groups together between sequencing technologies, and the six outbreak strains (Ec01-Ec06) are clustered together differentiating them from the remaining isolates (Ec07-Ec12) of the same ST type. This coincide with previous studies and the epidemiological data that concluded that the isolates Ec01-Ec06 are from the same outbreak, whereas the isolates Ec07-Ec12 were acquired from different sources in different foreign countries (6). Likewise, the clustering of Illumina and hac_m Q10 ONT data revealed the same clustering, where alignment trimming were performed with a minimum length of 10 between called SNPs (see Figure 3). The Newick files from the trees in Figure 1-3 can be seen in S2. The same topology was achieved using IQtree and FastTree on the three combinations of Illumina, fast base-called ONT Q8 and hac_m ONT Q10 data (see S3). This suggests that the alignments and trimming of alignments determine the clustering and phylogeny to a larger degree than the methods of clustering and phylogeny themselves. This coincides with the assumptions made by most maximum likelihood methods, which trust the alignments and are not build to differentiate between long closely related sequences, such as this study contains (20)(21). For distantly related sequences the maximum likelihood methods will produce more precise results, whereas the MINTyper pipeline will fail mostly due to the reference guided multiple alignment that does not account for large rearrangements of the genome.

**Figure 1.**
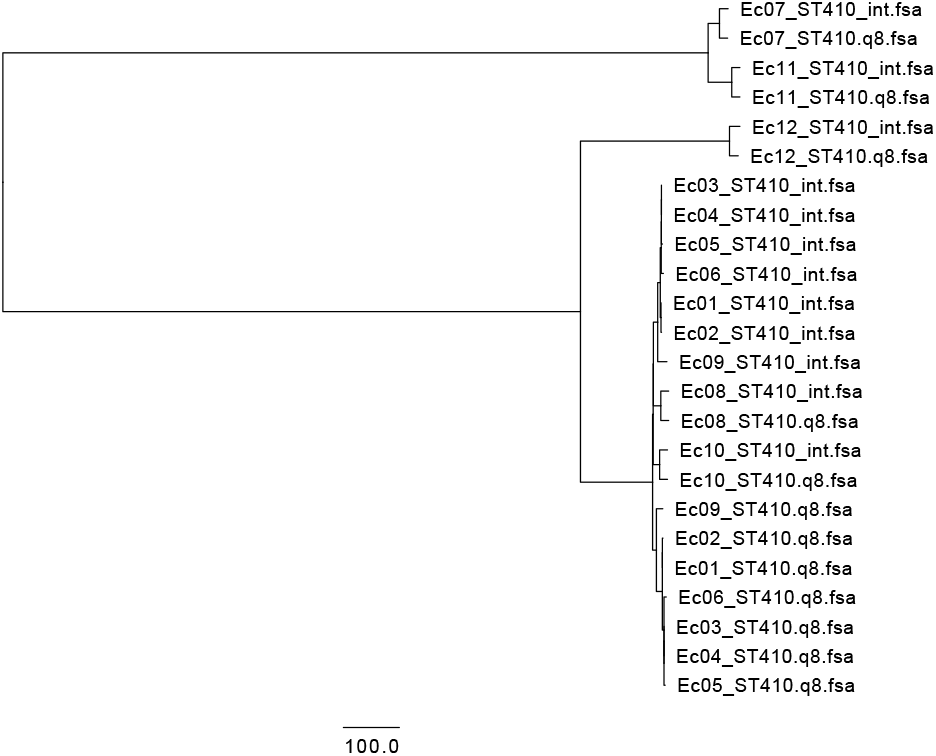
Clustering of sequences from Illumina (denoted int) and fast base-called Q8 ONT sequences (denoted Q8) of 12 *E. coli*, based on core genome SNPs without alignment trimming. Isolates Ec01-Ec06 are from an outbreak in Denmark, while Ec07-Ec12 originate from different foreign countries.

**Figure 2.**
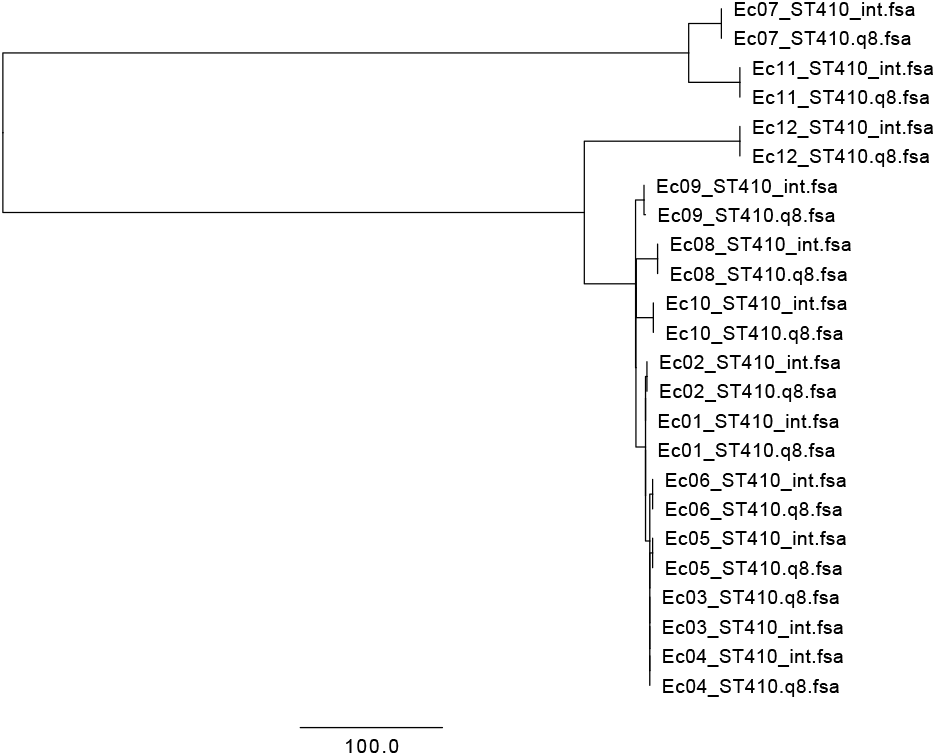
Clustering of sequences from Illumina (denoted int) and fast base-called Q8 ONT sequences (denoted Q8) of 12 *E. coli*, based on core genome SNPs. SNPs were trimmed away if they were within a proximity of 10, together with masking of DCM methylation-sites. Isolates Ec01-Ec06 are from an outbreak in Denmark, and Ec07-Ec12 originate from different foreign countries.

**Figure 3.**
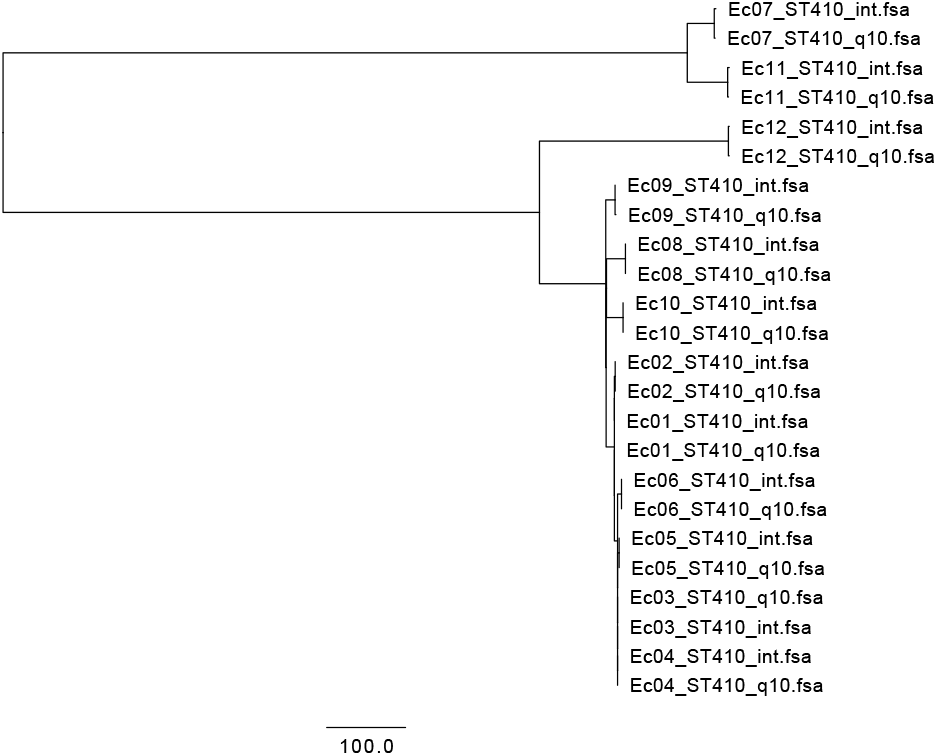
Clustering of sequences from Illumina (denoted int) and high-accuracy methylation-aware (hac_m) base-called Q10 ONT sequences (denoted Q10) of 12 *E. coli*, based on core genome SNPs. SNPs were trimmed away if they were within a proximity of 10. Isolates Ec01-Ec06 are from an outbreak in Denmark, and Ec07-Ec12 originate from different foreign countries.

### Loss of data

When performing either SNP or methylation-motif masking a certain part of the data is excluded from the analysis. Since the errors in the ONT MinION sequences are derived at the sequencing/base-calling stage, the best option to make a good clustering / phylogenetic analysis is to only look at the correctly sequenced parts of the isolates. In this study, we masked both insignificant base-calls (lower-case base-call letters), DCM motifs and SNPs in proximity of 10 bases of each other. When using MINTyper with no motif or proximity masking on fast Q8 MinION data, a total of 4183458/4767526 (87.7%) reference-genome bases were included in the distance matrix. When masking the DCM motifs and performing proximity masking on fast Q8 MinION data a total of 3835782/4767526 (80.5%) bases were included in the distance matrix.

Naturally, loosing data will lead to a less sensitive result, where true differences are at risk of being masked out. However, as was shown in Figure 1-3, we can produce more accurate results when trimming the alignments, even when it means loosing bases amounting to 8.3% of the included core-genome positions. Thus, in the case of SNP typing closely related isolates, it is more important to have a high quality of data rather than a high coverage of the core genome.

If we only employ masking of SNPs within a proximity of 10 and use the methylation-aware high-accuracy Q10 MinION data, we are able to include 4267999/4767526 (89.5%) bases in the analysis. As demonstrated in Figure 3, the structural errors found in Figure 1 no longer appears, and a greater fraction of base pairs are included in the analysis.

### Performance evaluation and comparison

To evaluate the performance of MINTypers ability to resolve outbreaks, we challenged MASH and CSIPhylogeny with the same combinations of Illumina and ONT data as MINTyper. MASH were tested with the sketch sizes: 1024, 1048576 and 4194304 in combination with a minimum threshold of *k*-mer count of: 1, 2, 8, 16 and 32. For all combinations of sketch size and minimum *k*-mer count, MASH were not able to cluster the outbreak sequences (Ec01-Ec06) together, but clustered mostly based on sequencing technology (see S4 and S5). CSIPhylogeny crashed when analysing both combinations of the ONT data after 4 hours with a peak memory of 256 GB. The crash were due to a malformed bam file, due to a flaw in the bam-format that cannot handle alignment cigars longer than 65535 operations.

In addition to the test above, MINTyper was tested with a Nanopore-only assembly as reference, using Unicycler v0.4.8-beta (minimap2 v2.17-r941, miniasm v0.3-v179, Racon 1.3.1) with default options (24) (25) (26) (27), and concurrent polishing using Medaka v1.2.1 (minimap2 v2.17-r941, samtools v1.10) with option “-m r941 min high g360” (25) (28), thus skipping the automatic reference-identification. Using the Nanopore-only assembly as reference revealed the same clusters as with the Illumina-Nanopore hybrid assembly of the “Escherichia coli strain AMA1167 chromosome, complete genome” reference. Nanopore-only assembly, distance matrices and Neighbor-Joining trees have been included in S6, S7 and S8 respectively. The computational requirements, in terms of CPU hours and peak memory, of the different methods are shown in table 2. All parts of the MINTyper pipeline can run multi-threaded, except for the automatic reference-identification. This can drastically lower the wall-time compared to the CPU time. The automatic reference-identification accounted for 3.3% to 7.6% of the total run time of MINTyper, and was the component responsible for the peak memory, except when FastTree was used to construct the trees. The alignments accounted for most of the CPU time used by MINTyper, where the Illumina samples had an average run time of 1.1 CPU-min., the fast base-called ONT Q8 used 7.9 CPU-min and the hac_m ONT Q10 used 5.3 CPU-min on average.

**Table 2.**
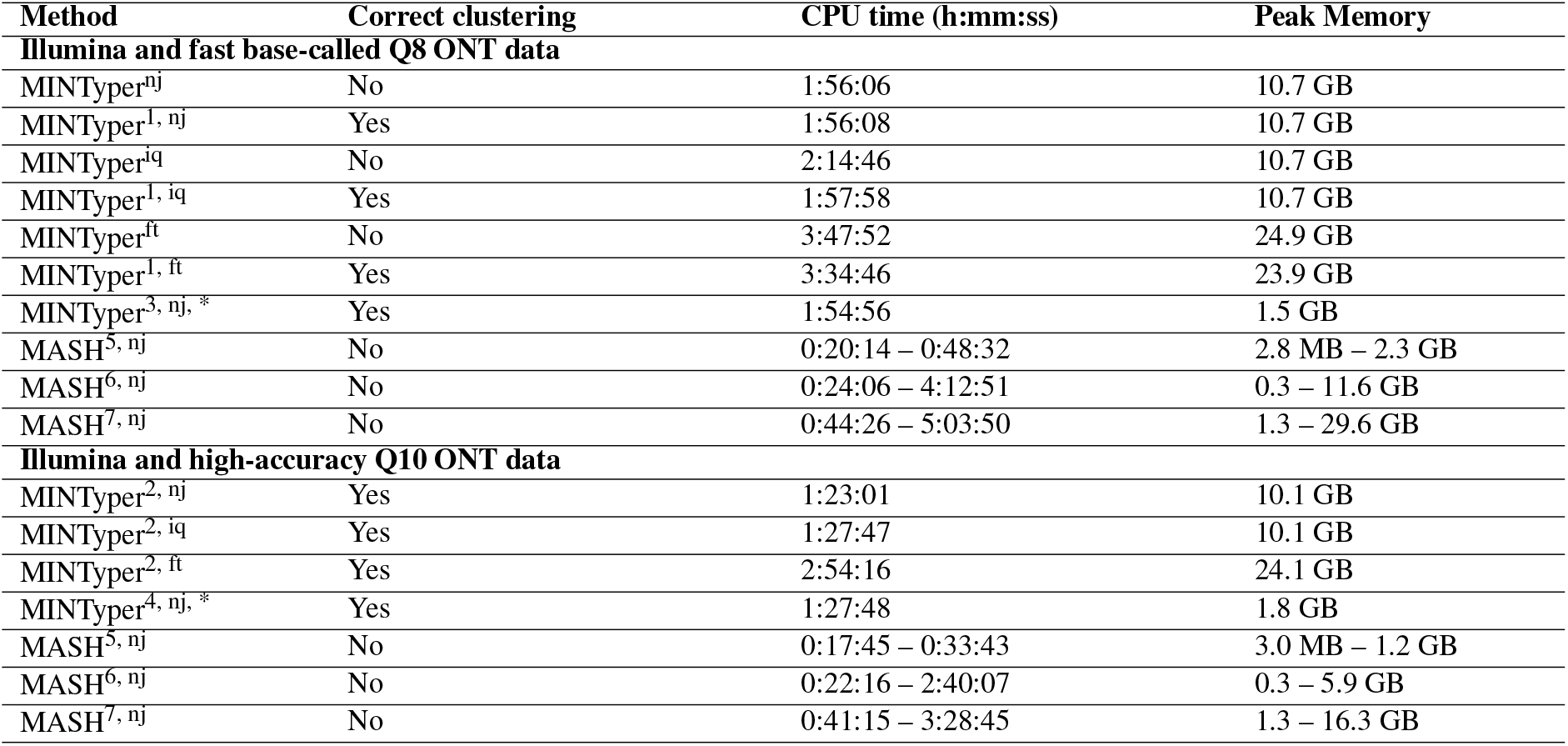
Computational requirements of tested methods against 12 E. coli isolates sequenced on Illumina and ONT MinIon with fast base-called Q8 and high-accuracy methylation-aware Q10 base-calling data. 1: Alignment trimming; Core-genome SNPs with a minimum distance of 10 between called SNPs and DCM-methylation masking, 3: Alignment trimming; Core-genome SNPs with a minimum distance of 10 between SNPs and DCM-methylation masking, 4: Alignment trimming; Core-genome SNPs with a minimum distance of 10 between SNPs, 5: Sketch size of 1024 with minimum k-mer occurrence thresholds varying from [1-32], 6: Sketch size of 1048576 with minimum k-mer occurrence thresholds varying from [1-32], 7: Sketch size of 4194304 with minimum k-mer occurrence thresholds varying from [1-32], nj: Neighbor-Joining tree-construction, iq: IQtree was used for tree-construction, ft: FastTree was used for tree-construction, *: Ec01 from the ONT data was assembled with Unicycler, polished with Medaka and used as reference.

| Method | Correct clustering | CPU time (h:mm:ss) | Peak Memory |
| --- | --- | --- | --- |
| **Illumina and fast base-called Q8 ONT data** |  |  |  |
| MINTyper<sup>nj</sup> | No | 1:56:06 | 10.7 GB |
| MINTyper<sup>1, nj</sup> | Yes | 1:56:08 | 10.7 GB |
| MINTyper<sup>iq</sup> | No | 2:14:46 | 10.7 GB |
| MINTyper<sup>1, iq</sup> | Yes | 1:57:58 | 10.7 GB |
| MINTyper<sup>ft</sup> | No | 3:47:52 | 24.9 GB |
| MINTyper<sup>1, ft</sup> | Yes | 3:34:46 | 23.9 GB |
| MINTyper<sup>3, nj, *</sup> | Yes | 1:54:56 | 1.5 GB |
| MASH<sup>5, nj</sup> | No | 0:20:14 – 0:48:32 | 2.8 MB – 2.3 GB |
| MASH<sup>6, nj</sup> | No | 0:24:06 – 4:12:51 | 0.3 – 11.6 GB |
| MASH<sup>7, nj</sup> | No | 0:44:26 – 5:03:50 | 1.3 – 29.6 GB |
| **Illumina and high-accuracy Q10 ONT data** |  |  |  |
| MINTyper<sup>2, nj</sup> | Yes | 1:23:01 | 10.1 GB |
| MINTyper<sup>2, iq</sup> | Yes | 1:27:47 | 10.1 GB |
| MINTyper<sup>2, ft</sup> | Yes | 2:54:16 | 24.1 GB |
| MINTyper<sup>4, nj, *</sup> | Yes | 1:27:48 | 1.8 GB |
| MASH<sup>5, nj</sup> | No | 0:17:45 – 0:33:43 | 3.0 MB – 1.2 GB |
| MASH<sup>6, nj</sup> | No | 0:22:16 – 2:40:07 | 0.3 – 5.9 GB |
| MASH<sup>7, nj</sup> | No | 0:41:15 – 3:28:45 | 1.3 – 16.3 GB |

## CONCLUSION

After performing separate experiments of MINTyper’s ability to cluster a set of 12 *E. coli* isolates with known relationships, it was found that ONT MinION long reads produced accurate clustering of the outbreak isolates with few discrepancies between sequencing technologies. This was achieved by employing KMA alignment, alignment trimming and DCM-methylation motif-masking. It was detected that in all 12 isolates the same systematic errors occurred in the MinION fast Q8 data. By masking the DCM motifs and trimming SNPs in close proximity all of these errors could be removed in 11 out of 12 samples, with only one SNP discrepancy in the remaining isolate. Running the analysis using methylation-aware high-accuracy Q10 MinION data, instead of fast Q8 data, lead to 1-3 SNP discrepancies in four samples.

Even though the trimming of alignments and the methylation-motif masking of fast Q8 ONT data resulted in a 8.3% data reduction, compared to no alignment trimming of the same data, the clonal clustering improved. When using ONT data of a higher quality, less alignment trimming were needed, and thus the methylation-motif masking could be excluded from the analysis. However, generating MinION data of a quality greater than fast Q8 can be extremely time consuming, and thus simply masking out error-generating motifs can be an effective tool when an urgent clustering is needed. Until the sequencing technology improves to allow for consistent, quick and precise sequencing and base-calling, MINTyper’s approach to apply long-read sequencing-data of lesser quality in outbreak detection has the potential to be a game changer in the field of genomic epidemiology.

### USAGE AND WEB-SERVICE

The source code for MINTyper are available at: https://github.com/MBHallgren/MINTyper.

A web-server service of MINTyper is available at: https://cge.cbs.dtu.dk/services/MINTyper/.

The data set used in this article was uploaded to ENA project accession no. PRJEB38543.

## ACKNOWLEDGEMENTS

We are grateful to Alfred F. Florensa and Judit Szarvas for assisting in the setup of the web-server and Frank Hansen and Karin Sixhøj Pedersen for excellent technical help in the laboratory.

## Supplementary

**S1**: Distances matrices for various trimming schemes. **S2**: Newick files of Figure 1-3. **S3**: Maximum likelihood trees from IQtree and FastTree. **S4**: MASH distance matrices. **S5**: Neighbour-Joining trees for S4. **S6**: Nanopore reference assemblies. **S7**: Distance matrices for S6. **S8**: Neighbour-Joining trees for S7.

